# Targeted gene deletion with *Sp*Cas9 and multiple guide RNAs in *Arabidopsis thaliana*: four are better than two

**DOI:** 10.1101/2023.01.10.523375

**Authors:** Jana Ordon, Niklas Kiel, Dieter Becker, Carola Kretschmer, Paul Schulze-Lefert, Johannes Stuttmann

**Affiliations:** Department of Plant-Microbe Interactions, Max-Planck Institute for Plant Breeding Research, D-50829 Cologne, Germany; Cluster of Excellence on Plant Sciences (CEPLAS), Max Planck Institute for Plant Breeding Research, Cologne, Germany; Institute for Biology, Department of Plant Genetics, Martin Luther University Halle-Wittenberg, D-06120 Halle, Germany; Institute for Biosafety in Plant Biotechnology, Federal Research Centre for Cultivated Plants, Julius Kühn-Institute (JKI), 06484 Quedlinburg, Germany; Aix Marseille University, CEA, CNRS, BIAM, UMR7265, LEMiRE (Rhizosphère et interactions sol-plante-microbiote), 13115 Saint-Paul lez Durance, France

## Abstract

**Background:** In plant genome editing, RNA-guided nucleases such as Cas9 from *Streptococcus pyogenes* (SpCas9) predominantly induce small insertions or deletions at target sites. This can be used for inactivation of protein-coding genes by frame shift mutations. However, in some cases, it may be advantageous to delete larger chromosomal segments. This is achieved by simultaneously inducing double strand breaks upstream and downstream of the fragment to be deleted. Experimental approaches for deletion induction have not been systematically evaluated.

**Results:** We designed three pairs of guide RNAs for deletion of the Arabidopsis *WRKY30* locus (~2.2 kb). We tested how the combination of guide RNA pairs and co-expression of the exonuclease TREX2 affect the frequency of *wrky30* deletions in editing experiments. Our data demonstrate that compared to one pair of guide RNAs, two pairs increase the frequency of chromosomal deletions. The exonuclease TREX2 enhanced mutation frequency at individual target sites and shifted the mutation profile towards larger deletions. However, TREX2 did not elevate the frequency of chromosomal deletions.

**Conclusions:** Multiplex editing with at least two pairs of guide RNAs (four guide RNAs in total) elevates the frequency of chromosomal deletions, and thus simplifies the selection of corresponding mutants. Co-expression of the TREX2 exonuclease can be used as a general strategy to increase editing efficiency in Arabidopsis without obvious negative effects.

## Background

Since the discovery of the mode of action of RNA-guided nucleases (RGNs; Gasiunas et al., 2012; Jinek et al., 2012; Cong et al., 2013; Mali et al., 2013), Cas9 from *Streptococcus pyogenes* (*Sp*Cas9) has become a routine tool for genome editing in many plant species. For mutagenesis of proteincoding genes, it is generally sufficient to program Cas9 for cleavage at a single target site within the gene of interest. Resulting double-strand breaks (DSBs) are mainly repaired by non-homologous end joining (NHEJ) in plant cells. As an error-prone process involving repeated RGN-mediated DNA cleavage upon precise repair, NHEJ frequently provokes small insertions or deletions at the initial DSB site. Indeed, +1/-1 nucleotide insertions/deletions (InDels) are the most frequently detected polymorphisms in CRISPR mutagenesis (Lemos et al., 2018; Chen et al., 2019). These small InDels provoke frame-shift mutations, which result in the disruption of protein-coding genes.

However, in a number of scenarios, it may be preferable to induce the deletion of a chromosomal segment. This may be the case, *e.g*., during mutagenesis of promoter regions to alter gene expression, functional interrogation of other non-coding sequences or deletion of gene clusters (Durr et al., 2018; Grutzner et al., 2021; Niu et al., 2021; Ordon et al., 2021; Wang et al., 2021). Also, remaining gene fragments may retain functionality, or the presence of alternative start codons (Bazykin and Kochetov, 2011) downstream of a target site or alternative splicing may lead to expression of a functional mRNA even after the introduction of small InDels. This can be prevented or excluded by the deletion of the full coding sequence. Further, deletions may help to characterize (supposedly) essential genes.

In site-specific mutagenesis with *Sp*Cas9 or other RGNs, bi-allelic mutations are often induced directly in primary transformants (Wang et al., 2015b; Grutzner et al., 2021). Thus, if an essential gene is targeted within the coding sequence, the majority of primary transformants will not survive. Obtaining informative material from such editing approaches requires discovery/isolation of primary transformants that are heterozygous for deleterious mutations, or that carry hypomorphic alleles. These two events are rare and cannot be specifically selected. *Sp*Cas9-induced chromosomal segment deletions often occur as hemizygous events in primary transformants and the second chromosomal copy may carry point mutations at *Sp*Cas9 target sites. Thus, if a deletion encompasses an essential gene (but point mutations at the target site do not affect gene function), hemizygous and viable primary transformants can be selected and further analyzed in a subsequent segregating population.

Chromosomal segment deletions (hereafter, deletions) are generated by inducing DSBs upstream and downstream of the targeted segment. In Arabidopsis (*Arabidopsis thaliana*), we have previously observed that deletion frequencies decreased with deletion size, and that point mutations at individual target sites were more frequent than deletions of the internal fragment (Ordon et al., 2017). Nonetheless, Arabidopsis lines carrying large deletions (e.g., > 40 - 80 kb) can be conveniently isolated from screening primary transformants by PCR, especially when using highly efficient nuclease systems (Grutzner et al., 2021). Also, in rice, large deletions occurred only in some transformants (Zhou et al., 2014), and point mutations at single target sites are more common (Pathak et al., 2019). In contrast, in tomato, deletions were more common than point mutations in at least one case, although a relatively small deletion (< 50 nt) was induced (Nekrasov et al., 2017).

A pair of guide RNAs programming Cas9 for cleavage at one target site upstream and one downstream of a given chromosomal segment is sufficient for the induction of deletions (dual targeting). However, addressing multiple up- and downstream target sites might increase the probability of losing the internal fragment and thus inducing the desired deletion. We therefore used two pairs of guide RNAs (four guide RNAs) when we intended to induce deletions in previous studies (Ordon et al., 2017; Ordon et al., 2020; Grutzner et al., 2021; Ordon et al., 2021), while others relied on dual targeting (e.g.; Wu et al., 2018). A systematic comparison to deduce design guidelines for deletion induction has not yet been conducted.

We compared here different constructs to evaluate whether increasing the number of guide RNA pairs or co-expression of a DNA exonuclease, TREX2, could enhance deletion frequencies in Arabidopsis. We chose the *WRKY30* locus for deletion. We show that increasing the number of guide RNAs from two to four enhanced deletion frequencies. In fact, we could detect bi-allelic deletions among primary transformants only when we edited with four guide RNAs. This facilitated the isolation of transgene-free mutants in the T_2_ generation without further screening. We confirmed mutant lines by long-read (PacBio) sequencing in the T_3_ generation. Co-expression of TREX2 exonuclease did not enhance the deletion frequency. However, TREX2 increased the frequency of point mutations at individual target sites two-fold and shifted the mutation spectrum towards larger deletions without adverse effects. Thus, co-expression of TREX2 can be used to augment mutation frequency during site-specific mutagenesis.

## Results

### Editing of the *WRKY30* locus in *Arabidopsis thaliana*

*We* aimed to generate an Arabidopsis *wrky30* mutant line by gene deletion, as we assumed *WRKY30* might be an essential gene (Scarpeci et al., 2013; Zou et al., 2019; Ma et al., 2021). Using this locus as a case study, we investigated whether increasing the number of guide RNA pairs and/or co-expression of the exonuclease TREX2 can enhance the frequency of deletions.

First, we assembled two new recipient vectors compatible with our <u>D</u>icot <u>G</u>enome <u>E</u>diting (pDGE) vector toolbox, pDGE1108 and pDGE1109 (Figure 1; Ordon et al., 2017; Stuttmann et al., 2021). These vectors contain a cassette for positive/negative selection by seed fluorescence (Fluorescence Accumulating Seed Technology (FAST); Shimada et al., 2010), the intron-optimized *zCas9i* gene under control of the *RPS5a* promoter (Tsutsui and Higashiyama, 2017; Ordon et al., 2020) and a “triple terminator” (t35S+tNbACT+Rb7-MAR; Diamos and Mason, 2018), and a *ccdB* cassette. The *ccdB* cassette is flanked by recognition sites for the Type IIs endonuclease *Bsa*I/*Eco31*I and can be replaced by one or multiple guide RNA transcriptional units by GoldenGate cloning to obtain final editing constructs (Figure 1a; Engler et al., 2008; Ordon et al., 2017).

**Figure 1:**
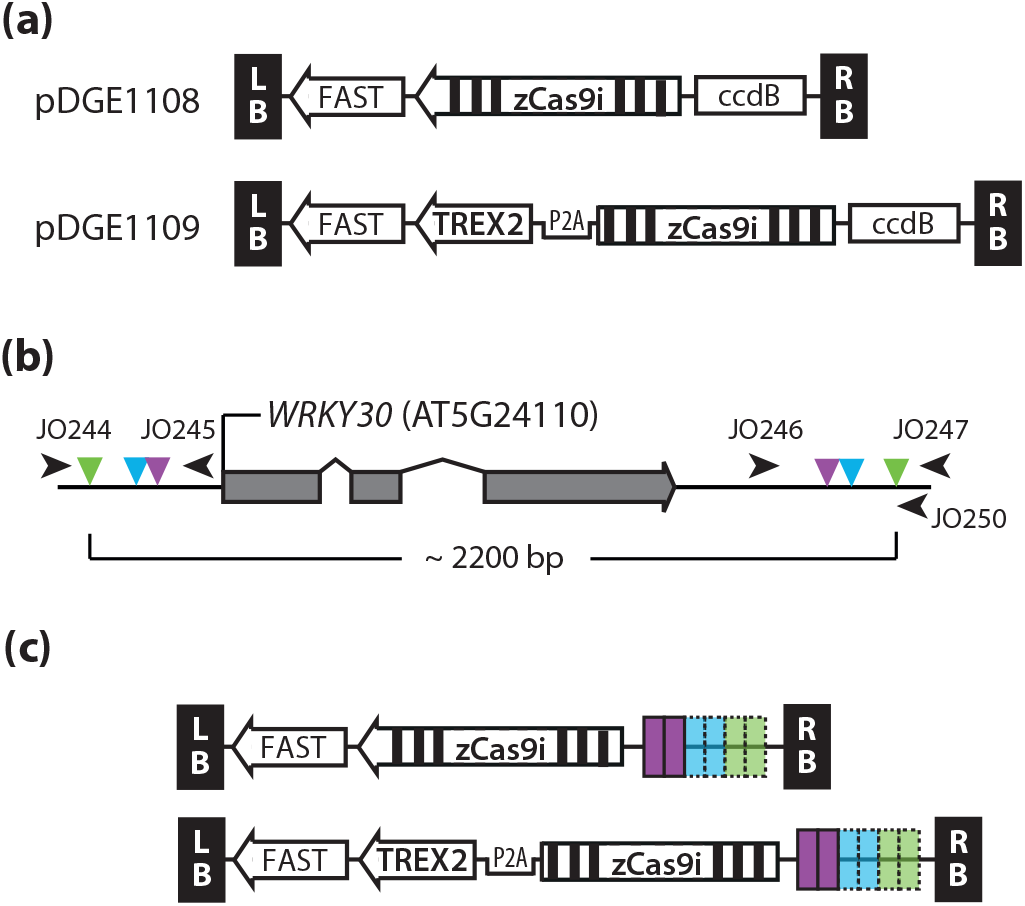
Constructs and target selection for deletion of the *AtWRKY30* locus. **(a)** Recipient vectors pDGE1108 and pDGE1109. The FAST (Fluorescence Accumulating Seed Technology) marker allows positive and negative selection of transgenics. The intron-optimized *zCas9i* gene is under control of the Arabidopsis *RPS5a* (*Ribosomal Particle S5a*) promoter and a chimeric terminator (“t-triple”: t35S-tNbACT-Rb7MAR; Diamos et al., 2018). **(b)** Scheme of the *WRKY30* locus and selected target sites (guide RNAs). Three pairs of target sites (inner, middle, outer) were selected, and respective guide RNAs designed. **(c)** Scheme of multiplex editing vectors assembled for *WRKY30* editing and containing or not *TREX2*. Constructs contain cassettes for expression of pairs of guide RNAs, pairs of pairs, or all six guide RNAs.

The chimeric “triple terminator” consists of the commonly used 35S terminator from Cauliflower Mosaic Virus fused to a terminator region from *Nicotiana benthamiana Actin3* and the Rb7 matrix attachment region from tobacco. In comparison to a construct containing only the *nopaline synthase* (*nos*) terminator, the triple terminator could previously increase the expression of GFP by up to 60-fold (Diamos and Mason, 2018). We analyzed Cas9 accumulation (35S promoter control) by agroinfiltration in *N. benthamiana* leaves. In comparison to the rbcS-E9 terminator (Wang et al., 2015b; Ordon et al., 2020), expression of Cas9 terminated by the chimeric triple terminator or by a tobacco extensin terminator (tNbEU; Diamos and Mason, 2018) resulted in a mild increase in protein accumulation (Figure S1).

In pDGE1109, the Cas9 expression cassette furthermore contains the *TREX2* coding sequence, fused 5’ to *zCas9i*, and separated by a P2A peptide-coding sequence (Figure 1a). When combined with a sequence-specific nuclease, exonucleases such as TREX2 promote end resection and thus augment mutagenesis frequency by increasing the error rate during NHEJ (Certo et al., 2012; Cermak et al., 2017). P2A mediates ribosomal skipping and thus the synthesis of TREX2 and Cas9 as individual polypeptides from a single mRNA (Donnelly et al., 2001; Wang et al., 2015a). When expressing *TREX2(P2A)-zCas9i* in agroinfiltration experiments, we did not detect additional higher molecular weight products using a Cas9-specific antibody (Figure S1). Further, protein amounts where similar with constructs containing *zCas9i* or *TREX2(P2A)-zCas9i* (Figure S1). Thus, TREX2 and Cas9 co-expression from a single transcript did not reduce Cas9 accumulation *in planta. We* conclude that read-through is not detectable for the used P2A sequence and that translation efficiency for the downstream polypeptide (Cas9) is not reduced.

To induce deletions of chromosomal fragments encompassing the *WRKY30* locus, we selected three pairs of target sites in flanking sequences (Figure 1b). The minimal/maximal distances between Cas9 cleavage sites were ~1,830 and 2,160 bp, respectively. The average gene size in Arabidopsis is approximately 2,200 bp (Derelle et al., 2006). The chosen deletion size is thus representative and applicable for deletion of many Arabidopsis genes. We designed guide RNAs corresponding to the selected target sites, and assembled constructs for expression of guide RNA pairs, pairs of guide RNA pairs, or all six guide RNAs, in both pDGE1108 and pDGE1109 (Figure 1c). This resulted in 14 different constructs (Table 1), which were transformed into Arabidopsis accession Col-0 by floral dipping.

**Table 1.**
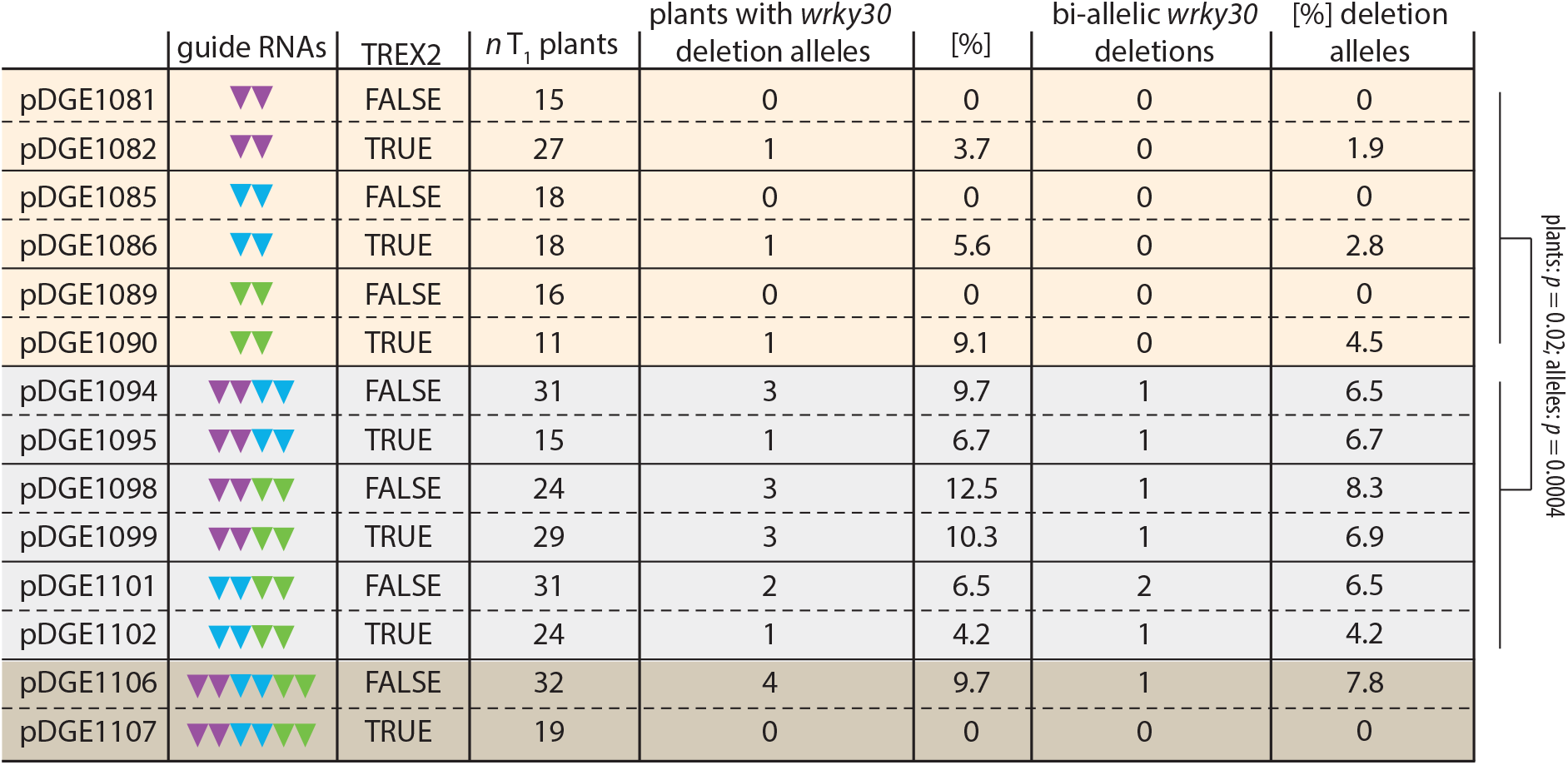

### Multiple cuts into flanking DNA genomic sequences increase deletion frequency

We assessed the frequency of gene deletions at the *WRKY30* locus by PCR screening in the T_1_ generation (Table 1, Figures S2-S4). Primary transformants were selected by seed fluorescence, and T_1_ plants (n = 11-32) genotyped using primers flanking the *WRKY30* locus and *Sp*Cas9 target sites (Figure 1b; JO244/247). Among transformants harboring constructs with guide RNA pairs (two guide RNAs), deletion alleles were detected at low frequencies. Bi-allelic deletions were not detected. Importantly, the frequency of deletion alleles increased when pairs of guide RNA pairs (4 guide RNAs) were expressed (Table S1). In this case, candidate bi-allelic deletion lines were recovered from all T_1_ populations. The frequency of deletion alleles was significantly elevated at the level of plants with deletion alleles (Student’s t-test, p = 0.02) and when comparing the absolute number of deletion alleles (p = 0.0004) between constructs with two or four guide RNAs. Deletion of *WRKY30* by six guide RNAs was detected in transformants expressing Cas9 (without TREX2) at frequencies similar to those obtained of the four guide RNAs. No deletion alleles were detected in transformants expressing six guide RNAs and TREX2-zCas9i (Table 1, Figure S4). No significant differences were detected when comparing plants with deletions or the absolute number of deletion alleles between populations with or without *TREX2* (Student’s t-test; p = 0.93 or p = 0.87, respectively).

Overall, we conclude that using four guide RNAs instead of two increases deletion frequency. Expression of six guide RNAs did not further elevate deletion frequency in our experiments; a result based on a limited number of observations. We did not detect a difference in deletion frequency with simultaneous expression of TREX2 and Cas9.

### Co-expression of the exonuclease TREX2 elevates mutation frequency and results in larger deletions

We further analyzed the effect of TREX2 and Cas9 co-expression at the level of individual target sites. We designed amplicons covering target sites up- and downstream of *WRKY30* (oligonucleotide combinations JO244/245 and JO246/247; Figure 1b). The respective amplicons were generated using DNA of pooled T_1_ individuals from transformation of pDGE1081-1090 (coding guide RNA pairs) by PCR, and subjected to amplicon deep sequencing. On average, approximately 10 % (6.1 – 13.7 %) of reads contained mutations at target sites when only Cas9 was expressed (Figure 2a,b). Mutation frequency was significantly elevated by TREX2 co-expression (p = 0.004, Student’s t-test) and increased on average two-fold (Figure 2a,b). Of note, although mutation frequency was improved with TREX2 in all comparisons, the effect size was variable and did not appear to correlate with the initial efficiency of a given guide RNA.

**Figure 2:**
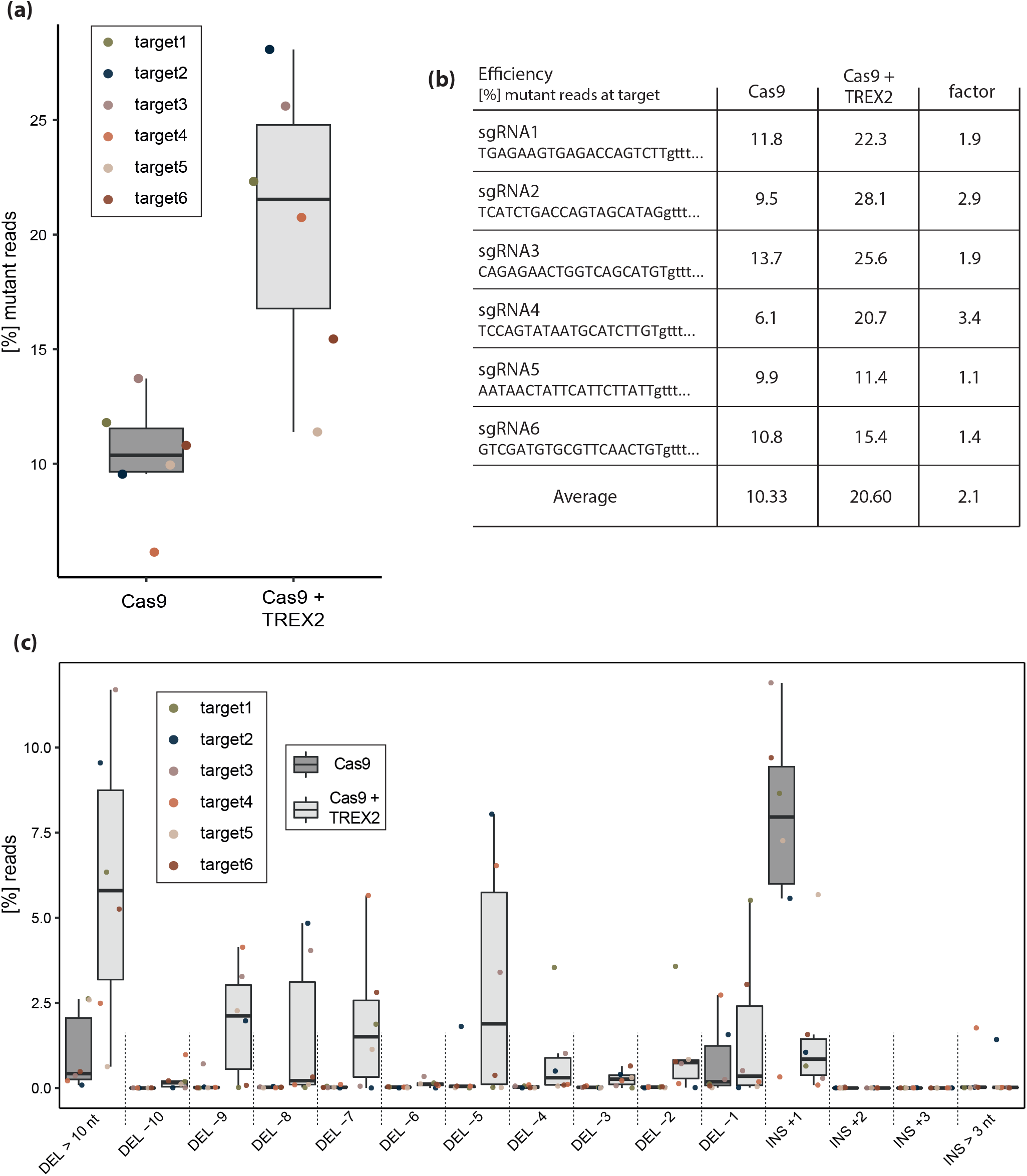
TREX2 improves editing efficiency and alters mutation profiles toward larger deletions. **(a)** Overall mutation frequencies at the six target sites with constructs containing or not *TREX2*. Amplicons containing the target sites up- or downstream of *WRKY30* were PCR-amplified using DNA from pooled T_1_ individuals (transformation of pDGE1081-1090). Up- and downstream amplicons for each construct were pooled and Illumina-sequenced. Mutation frequencies were determined using CRISPResso. **(b)** Guide RNA efficiencies at single target sites. As in a), but individual target sites +/- *TREX2* are shown. **(c)** Mutation profiles over six target sites upon editing with constructs containing or not the *TREX2* exonuclease gene. Samples and data analyses as in a), but frequencies of different InDels are illustrated. See also Figure S5 for individual target sites.

When editing with plain Cas9, insertions of a single nucleotide were detected most frequently (Figure 2c). Upon TREX2 co-expression, the frequency of insertions was significantly reduced and mutation profiles were shifted towards larger deletions (Figure 2c), as previously reported (Cermak et al., 2017; Weiss et al., 2020). Approximately 27 % of all InDel alleles were deletions of more than 10 nucleotides. Adverse effects of TREX2 co-expression, such as lowered numbers or reduced viability of primary transformants, were not observed.

In summary, we observed elevated mutation frequencies and a shift towards larger deletions at all target sites when TREX2 was co-expressed. On average, mutation frequencies doubled at target sites. TREX2 co-expression can thus be used as a general strategy to improve genome editing efficiencies in Arabidopsis.

### In-depth analysis of *wrky30* mutant lines and confirmation by long-read DNA sequencing

We selected T_2_ populations derived from four primary transformants for isolation of *wrky30* homozygous mutant lines. Putative bi-allelic deletions were detected in transformants 1094.29, 1101.12 and 1101.13 (Figure S3). Transformant 1098.5 was scored heterozygous for a deletion allele. We selected FAST-negative seeds from populations derived from these transformants to select against the presence of the T-DNA. T_2_ plants were sampled as pools of four plants (two pools per population), and pool DNA was used for genotyping (Figure S6). As with the primary transformants, bi-allelic and heterozygous deletion alleles were detected in the respective pools. Thus, mutations/deletions detected in the T_1_ generation were germline-transmitted to the T_2_ generation in all tested populations.

Single plants were propagated to the T_3_ generation, and PCR-genotyping was repeated on pools of five plants for lines with bi-allelic deletions (Figure 3a). Bi-allelic deletions were confirmed, and the lack of amplification of a *zCas9i*-specific PCR product confirmed the absence of the T-DNA.

**Figure 3:**
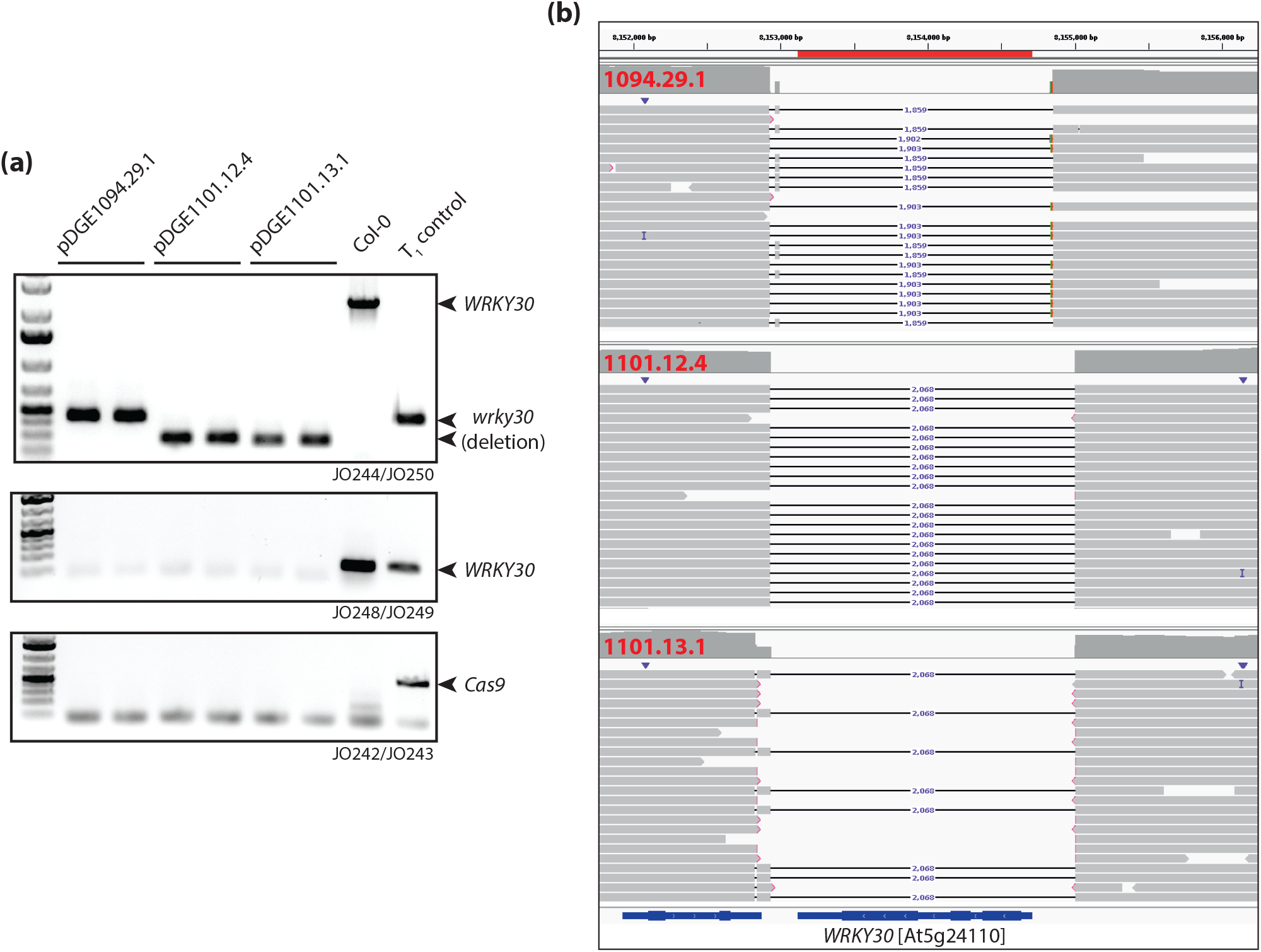
In-depth analysis of *wrky30* mutant lines by PCR and long-read DNA sequencing. **(a)** PCR genotyping of *wrky30* mutant lines. Pools were assembled from five randomly selected plants derived from indicated T_3_ populations; two pools per population. Corresponding DNAs were used for PCR genotyping (see Figure 1 for primer binding sites): Amplicon JO244/250 spans the *wrky30* locus; a smaller fragment is amplified in deletion lines. JO244/248 amplify a fragment within *WRKY30*, absent in deletion lines. JO242/243 amplify a fragment of the *zCas9i* gene to query presence/absence of the T-DNA. Col-0 and a T_1_ individual (1095.14) were included as controls. **(b)** Read mappings derived from long-read sequencing (Pacific Biosciences HiFi) of DNA pools (~ 20 plants) derived from indicated populations. Read mappings were visualized using Integrative Genome Viewer (Robinson et al., 2011). The *WRKY30* locus and ~ 1 kb of up-/downstream sequences is shown.

Pools of 20 plants per T_3_ population were used to extract DNA for long-read sequencing on a PacBio Sequel II system. HiFi reads were used for *de novo* assembly and contigs were aligned to TAIR10. Inspection of the *WRKY30* genomic region revealed that population 1094.29.1 was heterozygous for two different alleles: Deletions of approximately 1,860 bp between target sites of guide RNAs 3 and 5, and 1,900 bp between guide RNAs 3 and 4 (Figure 3b). An identical homozygous deletion of 2,068 bp between guide RNAs 1 and 4 was detected in populations 1101.12.4 and 1101.13.1 (Figure 3b). However, different mutations were detected at the target site of guide RNA6 in these lines, consistent agreement with independent lineages.

We used CRISPOR (Concordet and Haeussler, 2018) to predict possible off-targets of guide RNAs (Supplemental File S1). The respective locations were inspected in read mappings of PacBio data using IGV (Robinson et al., 2011). Mutations were not detected at any of the potential off-targeting sites.

Bi-allelic deletions were not detected among transformants expressing two guide RNAs. We observed a higher frequency of deletion alleles when expressing four guide RNAs. In this case, bi-allelic deletion mutants were detected in all populations tested. This enabled us to isolate clean *wrky30* deletion lines by selecting exclusively against the presence of the transgene.

## Discussion

The generation of deletions with RGNs such as *Sp*Cas9 requires simultaneous induction of DSBs up- and downstream of a targeted chromosomal segment (Figure 1). However, deletions are only induced in some individuals, as point mutations at single target sites are the more represented repair outcome (Ordon et al., 2017; Pathak et al., 2019). Here, we show that multiplexing with two guide RNA pairs (four guide RNAs; two target sites each up-/downstream) enhances the frequency of deletions compared to dual targeting (two guide RNAs; one target site up-/downstream). With most of the available RGN toolkits, the additional cost of integrating four, rather than two, guide RNAs into editing constructs is insignificant. Higher deletion frequencies may reduce the number of plants that need to be screened for isolation of candidate mutant lines. In our case, using four guide RNAs allowed us to directly isolate candidate lines with bi-allelic deletions in the T_1_ generation. Thus, only one selection against the transgene was required to obtain clean *wrky30* deletion lines. We conclude that four guide RNAs are better than two for inducing deletions.

RGN-induced deletions were reported, *e.g*., in Arabidopsis, soy bean, rice, tomato, *N. benthamiana* and *Catharanthus roseus* (Zhou et al., 2014; Nekrasov et al., 2017; Pathak et al., 2019; Duan et al., 2021; Grutzner et al., 2021). The frequency of deletions in primary T1 transformants is strongly locus/guide RNA-dependent (Ordon et al., 2017; Wu et al., 2018). For example, in *A. thaliana*, Wu et al. (2018) report a deletion frequency ranging from 5 % - 79 %, with deletions detected in 20 – 40% of primary transformants for most loci. In comparison, we obtained *wrky30* deletions at a moderate frequency of approximately 10 % (Table 1), but obtained deletions at higher frequencies (up to ~ 50%) at other loci with the same vector system (Grutzner et al., 2021; unpublished data). However, frequencies are expected to increase when editing at any locus with four guide RNAs. It should be noted that, previously, we observed inheritability of deletions with *RPS5a* promoter-driven *zCas9i* in six of eight tested lines in the T_2_ generation (Grutzner et al., 2021). In the present study, deletions were confirmed in T_2_ in all cases. So far, deletion alleles segregated at Mendelian ratios among non-transgenic T_2_ individuals in all populations tested. In contrast, Wu et al. (2018) recovered deletions at frequencies below 10 % for most analyzed T_2_ populations (1-90 %; Wu et al., 2018). Accordingly, corresponding primary transformants (*pcoCas9, Ubiquitin10* promoter control) were most likely somatic mosaics. This appears to be less of an issue with p*RPS5a*-driven *zCas9i*, or the differences might be due to tissues used for genotyping. We routinely use floral tissues for genotyping primary transformants. We assume that especially when bi-allelic deletions are detected in floral tissues, non-inheritability or low representation of deletion alleles in the T_2_ generation is highly unlikely.

Co-expression of TREX2 with *Sp*Cas9 resulted in a two-fold increase in the efficiency of point mutation generation (Figure 2). An approximately two-fold increase in mutation frequency was previously observed in tomato, barley and *Setaria viridis* protoplast experiments (Cermak et al., 2017; Weiss et al., 2020). Stable transgenic lines expressing TREX2 and Cas9, but not only Cas9, were reported for *S. viridis* (Weiss et al., 2020). Our direct comparison of efficiency at a total of six different target sites corroborate that TREX2 can robustly improve mutation frequency in stable transformants. Larger deletions obtained with TREX2 may also facilitate functional interrogation or inactivation of, *e.g*., small non-coding RNA genes or transcription factor-binding sites (Oliva et al., 2019).

TREX2 has also been reported to make genome editing at target sites even more precise: by avoiding repeated cleavage due to higher probability of error-prone repair, TREX2 reduces the number of translocations and large deletions that can occur, as rare events, at on-target sites (Yin et al., 2022). Yin et al. (2022) also reported that TREX2 outperformed several other tested exonucleases, and could not detect collateral damage activity. Consistent with this, we did not observe adverse effects of TREX2 co-expression in our stable Arabidopsis transgenic lines. Thus, it appears that TREX2 co-expression can be used as a general and robust strategy to elevate mutation frequency in genome editing.

We used whole genome re-sequencing with PacBio HiFi reads for verification of our mutant lines. In contrast to Sanger sequencing, this allowed us to not only determine the precise genotype at the *WRKY30* locus (Figure 3), but also to confirm the absence of off-target mutations or, e.g., translocations. At least for the moderate genome size of Arabidopsis, PacBio sequencing thus represents a cost- and labor-efficient approach for comprehensive verification of genome-edited lines.

We chose to delete the entire *WRKY30* gene rather than disrupt its coding sequence, as we supposed it could be essential (Scarpeci et al., 2013; Zou et al., 2019; Ma et al., 2021). The successful generation of bi-allelic *wrky30* deletion mutants – plants were indistinguishable from the wild type – demonstrates that this is not the case. However, as initially intended, hemizygous deletions were detected in multiple primary transformants (Figures S2-S4, Table 1). Segregation of the deletion allele was confirmed for one population (1098.5.; Figure S6). Accordingly, gene deletion can be used to generate material segregating for detrimental alleles in essential genes.

### Experimental outline for deletion induction with four guide RNAs in *Arabidopsis thaliana*

- Define chromosomal segment targeted for deletion.
- Select 200-300 bp of 5’ and 3’ flanking sequences for target site selection / guide RNA design. *E.g*., chop-chop (Labun et al., 2016) and CRISPOR (Concordet and Haeussler, 2018) are useful tools to scan for and evaluate target sites.
- Select two target sites each in 5’ and 3’ flanking sequences. Target sites should be offset to avoid steric hindrance among Cas9/guide RNA complexes. However, a large offset will lead to important differences between possible deletion outcomes, which may complicate PCR screening. We therefore recommend an offset of 50-100 bp between cleavage sites.
- Design corresponding guide RNAs, assemble construct for multiplex editing using available toolkits.
- Plant transformation. Logemann et al. (2006) provided a convenient protocol for Arabidopsis transformation by floral dipping.
- Select primary transformants by FAST or antibiotic/herbicide resistance. At least 30-40 primary transformants should be obtained.
- Design PCR primers: Flanking the desired deletion, and at least one internal oligonucleotide. Screen T_1_ transformants for occurrence of deletion alleles (flanking oligonucleotides) and presence of the targeted chromosomal fragment (one flanking and one internal oligonucleotide). We preferentially use floral tissues of T_1_ transformants for DNA extraction.
- Propagate ≥ 5 plants in which a deletion was detected to the T_2_ generation.
- Select against presence of the transgene: Non-fluorescent seeds when FAST is available.
- Repeat genotyping with T_2_ plants. Select homozygous. Confirm absence of Cas9 by PCR genotyping. Propagate selected plants to the T_3_ generation.
- Determine precise allele information by Sanger sequencing of PCR products or NGS using T_2_ or T_3_ material.

## Methods

### Plant growth conditions and transformation

*Arabidopsis thaliana* accession Columbia-0 (Col-0) plants were cultivated under short day conditions in a walk-in chamber (8h light, 23/21°C day/night, 60 % relative humidity) or in a greenhouse under long day conditions (16h light) for seed set. Arabidopsis was transformed by floral dipping as previously described (Logemann et al., 2006). Agrobacterium strain GV3101 pMP90 was used. Primary transformants (T_1_) were selected by seed fluorescence (Shimada et al., 2010) using a stereomicroscope equipped with an mCherry filter. Plants were grown in growth chambers for genotyping, and transferred to a “speed breeding chamber” (20h light) for seed production. In the T_2_ generation, non-fluorescent seeds were selected, respective plants genotyped and propagated to the next generation. *N. benthamiana* plants were cultivated in a greenhouse with a 16-h light period (sunlight and/or IP65 lamps [Philips] equipped with Agro 400 W bulbs [SON-T]; 130–150 μE m^-2^ s^-1^; switchpoint; 100 μE m^-2^ s^-1^), 60% relative humidity at 24/20°C (day/night).

### Molecular cloning and guide RNA design

The GoldenGate technique following the Modular Cloning syntax for hierarchical DNA assembly was used for clonings (Engler et al., 2008; Engler et al., 2014). Previously reported plasmids belonging to the Modular Cloning Toolkit and the MoClo Plant Parts I and II collections were used (Engler et al., 2014; Gantner et al., 2018). Recipient vectors pDGE1108 and pDGE1109 were assembled as previously described (Ordon et al., 2017; Stuttmann et al., 2021). A plasmid containing the TREX2 coding sequence was obtained via Addgene (#91026; Cermak et al., 2017). Oligonucleotides corresponding to the target sites TGAGAAGTGAGACCAGTCTTnGG (#1), TCATCTGACCAGTAGCATAGnGG (#2), CAGAGAACTGGTCAGCATGTnGG (#3), TCCAGTATAATGCATCTTGTnGG (#4), AATAACTATTCATTCTTATTnGG (#5), and GTCGATGTGCGTTCAACTGTnGG (#6) were cloned into guide RNA shuttle vectors containing the Arabidopsis U6-26 promoter described in Stuttmann et al. (2021). Final plant transformation vectors were generated by cloning guide RNA expression cassettes into pDGE1108/1109 as previously described (Stuttmann et al., 2021). Target sites were selected using CRISPOR (Concordet and Haeussler, 2018). Further details are provided in Supplemental File S2.

### Agroinfiltration and immunodetection

Four-to six-week-old *N. benthamiana* plants were used for agroinfiltration (OD_600_ = 0.4). Leaf discs were harvested three dpi, ground in liquid nitrogen and boiled in 2 x Laemmli buffer for protein extraction. Proteins were separated on 6 % SDS-PAA gels, and transferred to nitrocellulose membranes by tank blotting. Monoclonal antibody Abcam EPR18991 was used for detection of Cas9. HRP-conjugated secondary antibodies (GE Healthcare) were used. A mixture of SuperSignal West Pico and Femto was used for revelation on Kodak Biomax Light films.

### Genotyping, amplicon sequencing and data analysis

Oligonucleotides used for PCR genotyping are provided in Supplemental File S1. For initial deletion screening, a DNA purification-free PCR protocol was used (Jia et al., 2021). A standard CTAB protocol was used for further DNA extractions. For amplicon sequencing, PCR products were prepared on DNA of pooled T_1_ individuals using oligonucleotides JO244/245 and JO246/247, purified using a column kit, quantified on a Qbit, and sequenced by Genewiz (Amplicon-EZ). Data were analyzed using CRISPResso2 (Clement et al., 2019). Data contained in files “Indel_histogram” and “CRISPResso_quantification_of_editing_frequency” were used for preparation of figures 2 and S5.

### Long-read sequencing (PacBio)

Approximately 1 g of tissues derived from two week-old plants grown on 1/10 MS plates were used for DNA extraction. PacBio HiFi reads were filtered using BLASR (Chaisson and Tesler, 2012) to remove the PacBio 2kb sequence control. We employed a previously described approach for structural variant calling (Kim and Kim, 2022). We performed *de novo* assemblies using Flye 2.9-b1768 (Kolmogorov et al., 2020) with an estimated genome size of 135M and four polishing runs. We assessed the quality of the assembled contigs by (1) visualization with Bandage 0.8.1 (Wick et al., 2015), (2) calculation of the cumulative coverage and the N50 value as described (Kim and Kim, 2022), and (3) testing for the completeness of the assembly using BUSCO v5.2.2 in genome mode against the brassicales_odb10 database (Manni et al., 2021). Next, contigs were used for scaffolding with Ragtag v2.1.0 (Alonge et al., 2022) with default parameters and the TAIR10 Arabidopsis reference genome (https://ftp.ensemblgenomes.ebi.ac.uk/pub/plants/release-55/fasta/arabidopsis_thaliana/). To monitor the quality of the final assembly, we generated synteny plots of the final assembly using syri 1.6 (Goel et al., 2019) and plotsr 0.5.4 (Goel and Schneeberger, 2022). Structural variant calling was performed using SVIM-asm 1.0.2 (Heller and Vingron, 2020) in haploid mode with --max_sv_size set to 2500 in order to exclude larger structural variants as a consequence of mis-assemblies. Putative variants containing undefined nucleotides (N’s) as well as the centromeric regions were excluded from the analysis. The resulting variants were annotated using snpEff 4.3t (Cingolani et al., 2012) using the TAIR10 genome annotation as reference feature file. For IGV visualization, reads or assemblies were aligned against the TAIR10 reference genome using minimap2 2.24-r1122 (Li, 2021). The complete pipeline is implemented in Python 3.8.5 and depends on seaborn, pandas, biopython as well as bash sub-processes and is deposited on Github (https://github.com/bubu227/deletion-of-genes-with-SpCas9/blob/main/pacbio_analysis_pipeline.py).

## Acknowledgements

We are grateful to MLU & MPIPZ greenhouse staff for plant cultivation services. We acknowledge T. Cermak and colleagues for sharing the TREX2-containing plasmid pMOD_A0902 *via* Addgene. We thank the MPIPZ genome centre and Bruno Huettel for Arabidopsis genome resequencing. JS is grateful for support by the Julius Kuehn Institute.

## Author contributions

JS and JO designed experiments. JO, CK and DB performed experiments. JS and JO analyzed data and prepared figures. NK analyzed PacBio data. PSL supervised experiments. JS wrote the manuscripts. All authors read and approved the final manuscript.

## Data availability statement

All data is contained within the article or can be obtained through the authors. Plasmids pDGE1108 and pDGE1109 will be made available via Addgene. Pipeline for PacBio sequencing analysis is deposited on Github (https://github.com/bubu227/deletion-of-genes-with-SpCas9/blob/main/pacbio_analysis_pipeline.py).

## Supplemental Material

**Figure S1:** Expression of *zCas9i in N. benthamiana*.

**Figure S2:** T_1_ deletion screening upon editing with two guide RNAs.

**Figure S3:** T_1_ deletion screening upon editing with two guide RNAs.

**Figure S4:** T_1_ deletion screening upon editing with six guide RNAs.

**Figure S5:** Mutation (InDel) profiles in absence/presence of TREX2 at single target sites.

**Figure S6:** PCR-genotyping of putative *wrky30* deletion lines in the T_2_ generation.

**Supplemental File S1:** Potential off-targets predicted by CRISPOR.

**Supplemental File S2:** Plasmids, oligonucleotides, cloning details.

## References

Alonge, M., Lebeigle, L., Kirsche, M., Jenike, K., Ou, S., Aganezov, S., Wang, X., Lippman, Z.B., Schatz, M.C., and Soyk, S. (2022). Automated assembly scaffolding using RagTag elevates a new tomato system for high-throughput genome editing. Genome biology 23, 258.

Bazykin, G.A., and Kochetov, A.V. (2011). Alternative translation start sites are conserved in eukaryotic genomes. Nucleic Acids Res 39, 567–577.

Cermak, T., Curtin, S.J., Gil-Humanes, J., Cegan, R., Kono, T.J.Y., Konecna, E., Belanto, J.J., Starker, C. G., Mathre, J.W., Greenstein, R.L., and Voytas, D.F. (2017). A multi-purpose toolkit to enable advanced genome engineering in plants. The Plant cell.

Certo, M.T., Gwiazda, K.S., Kuhar, R., Sather, B., Curinga, G., Mandt, T., Brault, M., Lambert, A.R., Baxter, S.K., Jacoby, K., Ryu, B.Y., Kiem, H.P., Gouble, A., Paques, F., Rawlings, D.J., and Scharenberg, A.M. (2012). Coupling endonucleases with DNA end-processing enzymes to drive gene disruption. Nat Methods 9, 973–975.

Chaisson, M.J., and Tesler, G. (2012). Mapping single molecule sequencing reads using basic local alignment with successive refinement (BLASR): application and theory. BMC Bioinformatics 13, 238.

Chen, W., McKenna, A., Schreiber, J., Haeussler, M., Yin, Y., Agarwal, V., Noble, W.S., and Shendure, J. (2019). Massively parallel profiling and predictive modeling of the outcomes of CRISPR/Cas9-mediated double-strand break repair. Nucleic Acids Res 47, 7989–8003.

Cingolani, P., Platts, A., Wang le, L., Coon, M., Nguyen, T., Wang, L., Land, S.J., Lu, X., and Ruden, D. M. (2012). A program for annotating and predicting the effects of single nucleotide polymorphisms, SnpEff: SNPs in the genome of Drosophila melanogaster strain w1118; iso-2; iso-3. Fly (Austin) 6, 80–92.

Clement, K., Rees, H., Canver, M.C., Gehrke, J.M., Farouni, R., Hsu, J.Y., Cole, M.A., Liu, D.R., Joung, J.K., Bauer, D.E., and Pinello, L. (2019). CRISPResso2 provides accurate and rapid genome editing sequence analysis. Nature biotechnology 37, 224–226.

Concordet, J.P., and Haeussler, M. (2018). CRISPOR: intuitive guide selection for CRISPR/Cas9 genome editing experiments and screens. Nucleic Acids Res 46, W242–W245.

Cong, L., Ran, F.A., Cox, D., Lin, S., Barretto, R., Habib, N., Hsu, P.D., Wu, X., Jiang, W., Marraffini, L.A., and Zhang, F. (2013). Multiplex genome engineering using CRISPR/Cas systems. Science (New York, N.Y 339, 819–823.

Derelle, E., Ferraz, C., Rombauts, S., Rouze, P., Worden, A.Z., Robbens, S., Partensky, F., Degroeve, S., Echeynie, S., Cooke, R., Saeys, Y., Wuyts, J., Jabbari, K., Bowler, C., Panaud, O., Piegu, B., Ball, S.G., Ral, J.P., Bouget, F.Y., Piganeau, G., De Baets, B., Picard, A., Delseny, M., Demaille, J., Van de Peer, Y., and Moreau, H. (2006). Genome analysis of the smallest free-living eukaryote Ostreococcus tauri unveils many unique features. Proceedings of the National Academy of Sciences of the United States of America 103, 11647–11652.

Diamos, A.G., and Mason, H.S. (2018). Chimeric 3’ flanking regions strongly enhance gene expression in plants. Plant Biotechnol J 16, 1971–1982.

Donnelly, M.L.L., Luke, G., Mehrotra, A., Li, X., Hughes, L.E., Gani, D., and Ryan, M.D. (2001). Analysis of the aphthovirus 2A/2B polyprotein ‘cleavage’ mechanism indicates not a proteolytic reaction, but a novel translational effect: a putative ribosomal ‘skip’. J Gen Virol 82, 1013–1025.

Duan, K., Cheng, Y., Ji, J., Wang, C., Wei, Y., and Wang, Y. (2021). Large chromosomal segment deletions by CRISPR/LbCpf1-mediated multiplex gene editing in soybean. J Integr Plant Biol 63, 1620–1631.

Durr, J., Papareddy, R., Nakajima, K., and Gutierrez-Marcos, J. (2018). Highly efficient heritable targeted deletions of gene clusters and non-coding regulatory regions in Arabidopsis using CRISPR/Cas9. Sci Rep 8, 4443.

Engler, C., Kandzia, R., and Marillonnet, S. (2008). A one pot, one step, precision cloning method with high throughput capability. PLoS ONE 3, e3647.

Engler, C., Youles, M., Gruetzner, R., Ehnert, T.M., Werner, S., Jones, J.D., Patron, N.J., and Marillonnet, S. (2014). A Golden Gate Modular Cloning Toolbox for Plants. ACS synthetic biology.

Gantner, J., Ordon, J., Ilse, T., Kretschmer, C., Gruetzner, R., Lofke, C., Dagdas, Y., Burstenbinder, K., Marillonnet, S., and Stuttmann, J. (2018). Peripheral infrastructure vectors and an extended set of plant parts for the Modular Cloning system. PLoS ONE 13, e0197185.

Gasiunas, G., Barrangou, R., Horvath, P., and Siksnys, V. (2012). Cas9-crRNA ribonucleoprotein complex mediates specific DNA cleavage for adaptive immunity in bacteria. Proceedings of the National Academy of Sciences of the United States of America 109, E2579–2586.

Goel, M., and Schneeberger, K. (2022). plotsr: visualizing structural similarities and rearrangements between multiple genomes. Bioinformatics 38, 2922–2926.

Goel, M., Sun, H., Jiao, W.B., and Schneeberger, K. (2019). SyRI: finding genomic rearrangements and local sequence differences from whole-genome assemblies. Genome biology 20, 277.

Grutzner, R., Martin, P., Horn, C., Mortensen, S., Cram, E.J., Lee-Parsons, C.W.T., Stuttmann, J., and Marillonnet, S. (2021). High-efficiency genome editing in plants mediated by a Cas9 gene containing multiple introns. Plant Commun 2, 100135.

Heller, D., and Vingron, M. (2020). SVIM-asm: Structural variant detection from haploid and diploid genome assemblies. Bioinformatics 36, 5519–5521.

Jia, Z., Han, X., and Tsuda, K. (2021). An Efficient Method for DNA Purification-Free PCR from Plant Tissue. Curr Protoc 1, e289.

Jinek, M., Chylinski, K., Fonfara, I., Hauer, M., Doudna, J.A., and Charpentier, E. (2012). A programmable dual-RNA-guided DNA endonuclease in adaptive bacterial immunity. Science (New York, N.Y 337, 816–821.

Kim, J., and Kim, C. (2022). A beginner’s guide to assembling a draft genome and analyzing structural variants with long-read sequencing technologies. STAR Protoc 3, 101506.

Kolmogorov, M., Bickhart, D.M., Behsaz, B., Gurevich, A., Rayko, M., Shin, S.B., Kuhn, K., Yuan, J., Polevikov, E., Smith, T.P.L., and Pevzner, P.A. (2020). metaFlye: scalable long-read metagenome assembly using repeat graphs. Nat Methods 17, 1103–1110.

Labun, K., Montague, T.G., Gagnon, J.A., Thyme, S.B., and Valen, E. (2016). CHOPCHOP v2: a web tool for the next generation of CRISPR genome engineering. Nucleic Acids Res 44, W272–276.

Lemos, B.R., Kaplan, A.C., Bae, J.E., Ferrazzoli, A.E., Kuo, J., Anand, R.P., Waterman, D.P., and Haber, J.E. (2018). CRISPR/Cas9 cleavages in budding yeast reveal templated insertions and strand-specific insertion/deletion profiles. Proceedings of the National Academy of Sciences of the United States of America 115, E2040–E2047.

Li, H. (2021). New strategies to improve minimap2 alignment accuracy. Bioinformatics 37, 4572–4574.

Logemann, E., Birkenbihl, R.P., Ulker, B., and Somssich, I.E. (2006). An improved method for preparing Agrobacterium cells that simplifies the Arabidopsis transformation protocol. Plant methods 2, 16.

Ma, K.W., Niu, Y., Jia, Y., Ordon, J., Copeland, C., Emonet, A., Geldner, N., Guan, R., Stolze, S.C., Nakagami, H., Garrido-Oter, R., and Schulze-Lefert, P. (2021). Coordination of microbe-host homeostasis by crosstalk with plant innate immunity. Nat Plants 7, 814–825.

Mali, P., Yang, L., Esvelt, K.M., Aach, J., Guell, M., DiCarlo, J.E., Norville, J.E., and Church, G.M. (2013). RNA-guided human genome engineering via Cas9. Science (New York, N.Y 339, 823–826.

Manni, M., Berkeley, M.R., Seppey, M., Simao, F.A., and Zdobnov, E.M. (2021). BUSCO Update: Novel and Streamlined Workflows along with Broader and Deeper Phylogenetic Coverage for Scoring of Eukaryotic, Prokaryotic, and Viral Genomes. Mol Biol Evol 38, 4647–4654.

Nekrasov, V., Wang, C., Win, J., Lanz, C., Weigel, D., and Kamoun, S. (2017). Rapid generation of a transgene-free powdery mildew resistant tomato by genome deletion. Sci Rep 7, 482.

Niu, F., Jiang, Q., Sun, X., Hu, Z., Wang, L., and Zhang, H. (2021). Large DNA fragment deletion in lncRNA77580 regulates neighboring gene expression in soybean (Glycine max). Funct Plant Biol 48, 1139–1147.

Oliva, R., Ji, C., Atienza-Grande, G., Huguet-Tapia, J.C., Perez-Quintero, A., Li, T., Eom, J.S., Li, C., Nguyen, H., Liu, B., Auguy, F., Sciallano, C., Luu, V.T., Dossa, G.S., Cunnac, S., Schmidt, S.M., Slamet-Loedin, I.H., Vera Cruz, C., Szurek, B., Frommer, W.B., White, F.F., and Yang, B. (2019). Broad-spectrum resistance to bacterial blight in rice using genome editing. Nature biotechnology 37, 1344–1350.

Ordon, J., Bressan, M., Kretschmer, C., Dall’Osto, L., Marillonnet, S., Bassi, R., and Stuttmann, J. (2020). Optimized Cas9 expression systems for highly efficient Arabidopsis genome editing facilitate isolation of complex alleles in a single generation. Functional & integrative genomics 20, 151–162.

Ordon, J., Martin, P., Erickson, J.L., Ferik, F., Balcke, G., Bonas, U., and Stuttmann, J. (2021). Disentangling cause and consequence: genetic dissection of the DANGEROUS MIX2 risk locus, and activation of the DM2h NLR in autoimmunity. Plant J 106, 1008–1023.

Ordon, J., Gantner, J., Kemna, J., Schwalgun, L., Reschke, M., Streubel, J., Boch, J., and Stuttmann, J. (2017). Generation of chromosomal deletions in dicotyledonous plants employing a userfriendly genome editing toolkit. Plant J 89, 155–168.

Pathak, B., Zhao, S., Manoharan, M., and Srivastava, V. (2019). Dual-targeting by CRISPR/Cas9 leads to efficient point mutagenesis but only rare targeted deletions in the rice genome. 3 Biotech 9, 158.

Robinson, J.T., Thorvaldsdottir, H., Winckler, W., Guttman, M., Lander, E.S., Getz, G., and Mesirov, J.P. (2011). Integrative genomics viewer. Nature biotechnology 29, 24–26.

Scarpeci, T.E., Zanor, M.I., Mueller-Roeber, B., and Valle, E.M. (2013). Overexpression of AtWRKY30 enhances abiotic stress tolerance during early growth stages in Arabidopsis thaliana. Plant molecular biology 83, 265–277.

Shimada, T.L., Shimada, T., and Hara-Nishimura, I. (2010). A rapid and non-destructive screenable marker, FAST, for identifying transformed seeds of Arabidopsis thaliana. Plant J 61, 519–528.

Stuttmann, J., Barthel, K., Martin, P., Ordon, J., Erickson, J.L., Herr, R., Ferik, F., Kretschmer, C., Berner, T., Keilwagen, J., Marillonnet, S., and Bonas, U. (2021). Highly efficient multiplex editing: one-shot generation of 8x Nicotiana benthamiana and 12x Arabidopsis mutants. Plant J 106, 8–22.

Tsutsui, H., and Higashiyama, T. (2017). pKAMA-ITACHI Vectors for Highly Efficient CRISPR/Cas9-Mediated Gene Knockout in Arabidopsis thaliana. Plant Cell Physiol 58, 46–56.

Wang, X., Aguirre, L., Rodriguez-Leal, D., Hendelman, A., Benoit, M., and Lippman, Z.B. (2021). Dissecting cis-regulatory control of quantitative trait variation in a plant stem cell circuit. Nat Plants 7, 419–427.

Wang, Y., Wang, F., Wang, R., Zhao, P., and Xia, Q. (2015a). 2A self-cleaving peptide-based multi-gene expression system in the silkworm Bombyx mori. Sci Rep 5, 16273.

Wang, Z.P., Xing, H.L., Dong, L., Zhang, H.Y., Han, C.Y., Wang, X.C., and Chen, Q.J. (2015b). Egg cell-specific promoter-controlled CRISPR/Cas9 efficiently generates homozygous mutants for multiple target genes in Arabidopsis in a single generation. Genome biology 16, 144.

Weiss, T., Wang, C., Kang, X., Zhao, H., Elena Gamo, M., Starker, C.G., Crisp, P.A., Zhou, P., Springer, N.M., Voytas, D.F., and Zhang, F. (2020). Optimization of multiplexed CRISPR/Cas9 system for highly efficient genome editing in Setaria viridis. Plant J 104, 828–838.

Wick, R.R., Schultz, M.B., Zobel, J., and Holt, K.E. (2015). Bandage: interactive visualization of de novo genome assemblies. Bioinformatics 31, 3350–3352.

Wu, R., Lucke, M., Jang, Y.T., Zhu, W., Symeonidi, E., Wang, C., Fitz, J., Xi, W., Schwab, R., and Weigel, D. (2018). An efficient CRISPR vector toolbox for engineering large deletions in Arabidopsis thaliana. Plant methods 14, 65.

Yin, J., Lu, R., Xin, C., Wang, Y., Ling, X., Li, D., Zhang, W., Liu, M., Xie, W., Kong, L., Si, W., Wei, P., Xiao, B., Lee, H.Y., Liu, T., and Hu, J. (2022). Cas9 exo-endonuclease eliminates chromosomal translocations during genome editing. Nat Commun 13, 1204.

Zhou, H., Liu, B., Weeks, D.P., Spalding, M.H., and Yang, B. (2014). Large chromosomal deletions and heritable small genetic changes induced by CRISPR/Cas9 in rice. Nucleic Acids Res 42, 10903–10914.

Zou, L., Yang, F., Ma, Y., Wu, Q., Yi, K., and Zhang, D. (2019). Transcription factor WRKY30 mediates resistance to Cucumber mosaic virus in Arabidopsis. Biochemical and biophysical research communications 517, 118–124.

