## Supplementary material for "Targeted gene deletion with *Sp*Cas9 and multiple guide RNAs in *Arabidopsis thaliana*: four are better than two": Figure S1

(a)

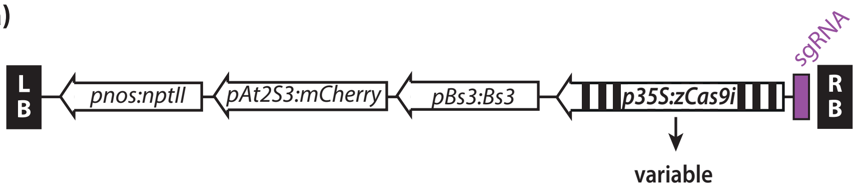


(b)

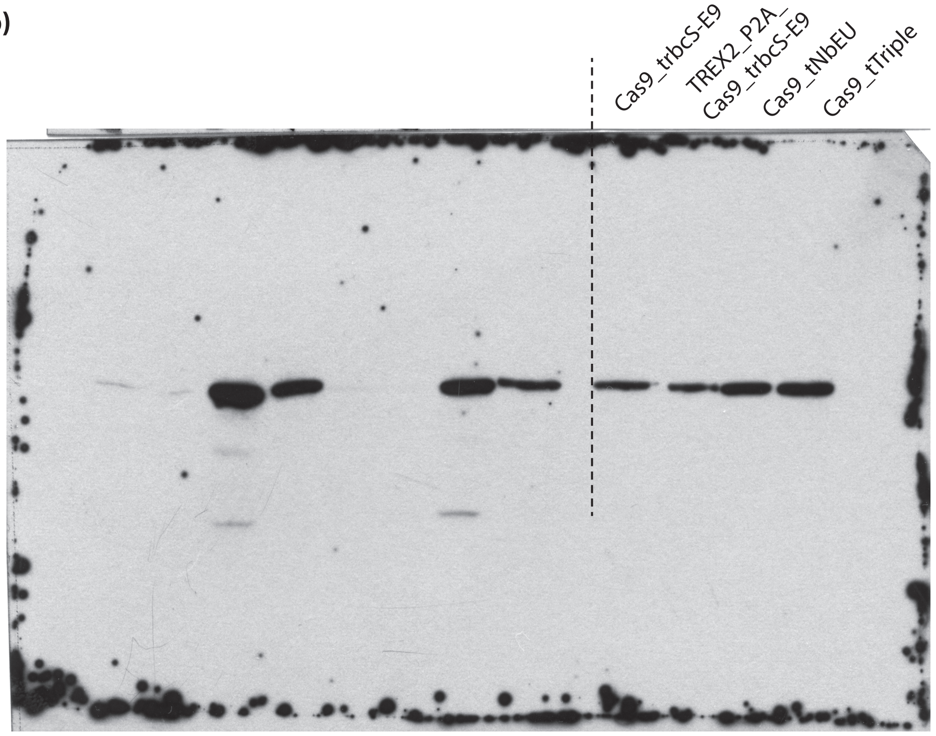


**Figure S1:** Expression of *zCas9i* in *N. benthamiana.*

**a)** Scheme of constructs used for expression in *N. benthamiana*. All constructs contained kanamycin-resistance as plant selectable marker, two cassettes designed for counter-selection (described in Stuttmann et al., 2021), an sgRNA expression unit and a cassette for zCas9i expression. Only the cassette for zCas9i expression was variable between constructs. zCas9i was expressed under 35S promoter control.

**b)** Immunodetection of Cas9. Agrobacterium strains for expression of Cas9 from the indicated cassettes were used for agroinfiltration at $OD_{600}$=0.4. Leaf discs were harvested three days post infiltration, ground in liquid nitrogen and boiled in Laemmli buffer for protein extraction. Proteins were seperated on a 6 % PAA gel and transferred to a nitrocellulose membrane. Cas9 was detected using a monoclonal $\alpha$-Cas9 antibody (Abcam EPR18991) and a HRP-coupled secondary antibody. Leaf discs originating from three independent replicates of the experiment were pooled for protein extraction. The entire membrane is shown to illustrate the absence of a signal corresponding to TREX2-Cas9 and/or degradation products. Left halve of the membrane corresponds to Cas9 detection in unrelated samples; it was included to clarify membrane margins.
