## Supplementary material for "Targeted gene deletion with *Sp*Cas9 and multiple guide RNAs in *Arabidopsis thaliana*: four are better than two": Figure S3

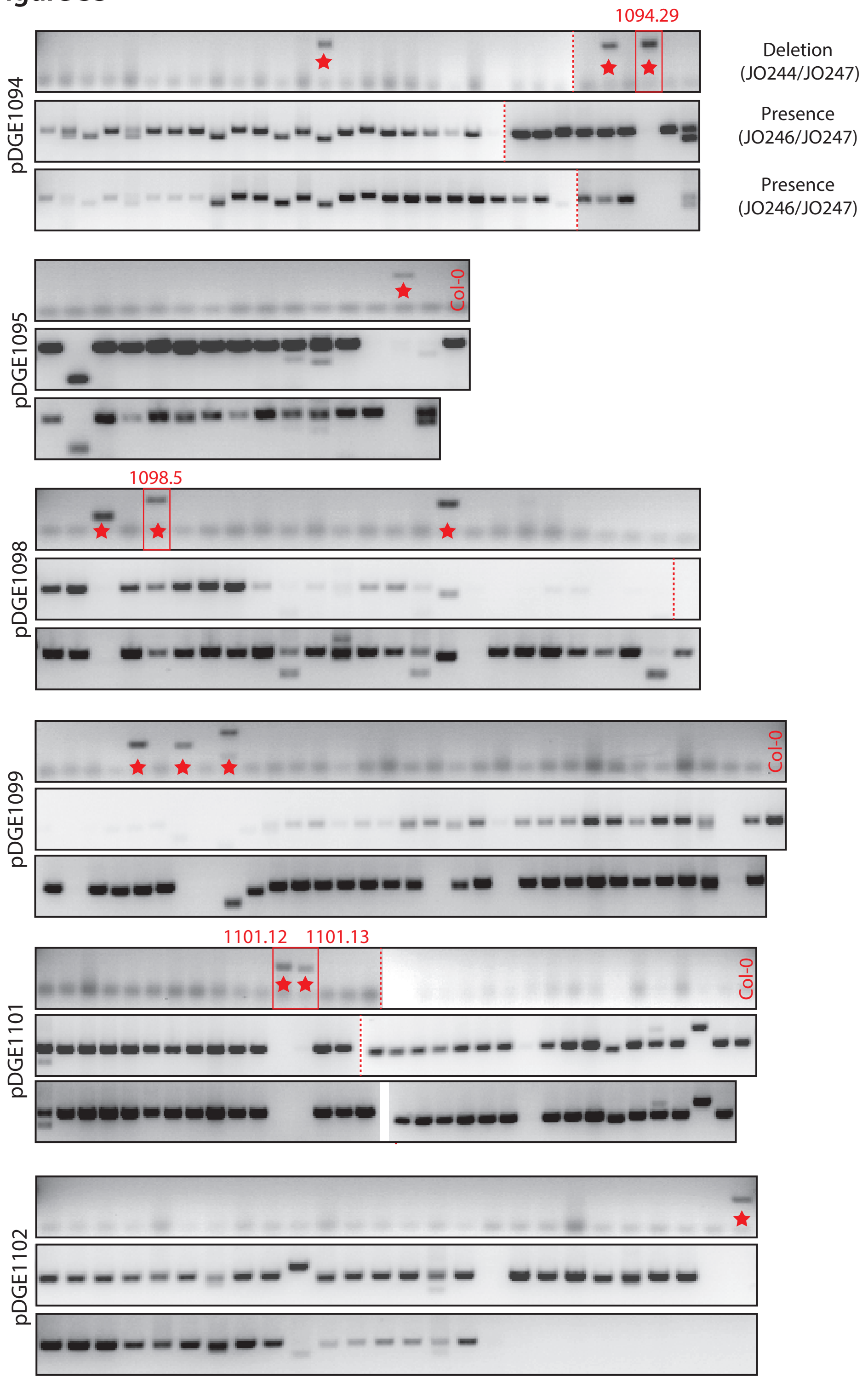


**Figure S3:** $T_1$ deletion screening upon editing with four guide RNAs.

Primary transformants ($T_1$) from transformation of indicated constructs were screened, by PCR, for presence of a large deletion encompassing the *WRKY30* locus (top PCR, oligonucleotides JO244/247). A second amplicon (oligonucleotides JO246/247) queries presence of the *WRKY30* locus, and serves as a control for DNA quality. PCR signals scored as presence of a deletion allele are marked with a star. Individuals for which neither PCR produced a signal were not counted. Grey boxes mask gel areas that were not considered, and dashed lines mark boundaries of spliced images. PCR signals corresponding to plants selected for further analysis in the $T_2$ generation are boxed and numbered (red).
