## Supplementary material for "Targeted gene deletion with *Sp*Cas9 and multiple guide RNAs in *Arabidopsis thaliana*: four are better than two": Figure S4

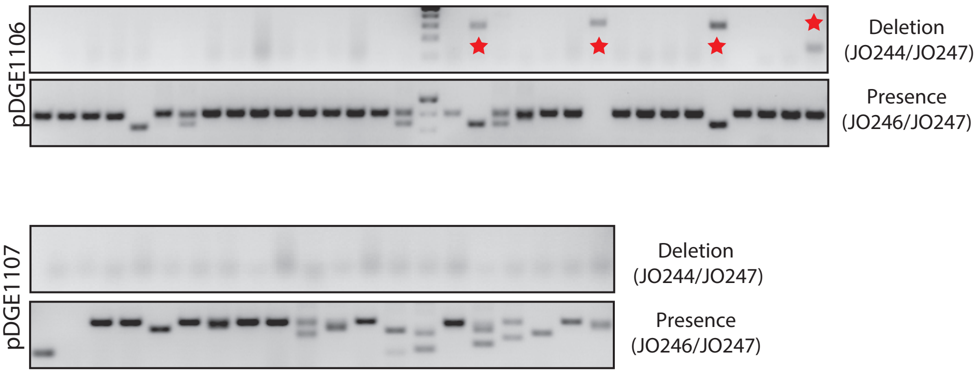


**Figure S4:** $T_1$ deletion screening upon editing with six guide RNAs.

Primary transformants ($T_1$) from transformation of indicated constructs were screened, by PCR, for presence of a large deletion encompassing the *WRKY30* locus (top PCR, oligonucleotides JO244/247). A second amplicon (oligonucleotides JO246/247) queries presence of the *WRKY30* locus, and serves as a control for DNA quality. PCR signals scored as presence of a deletion allele are marked with a star. Individuals for which neither PCR produced a signal were not counted. Grey boxes mask gel areas that were not considered, and dashed lines mark boundaries of spliced images.
