## Supplementary material for "Targeted gene deletion with *Sp*Cas9 and multiple guide RNAs in *Arabidopsis thaliana*: four are better than two": Figure S5

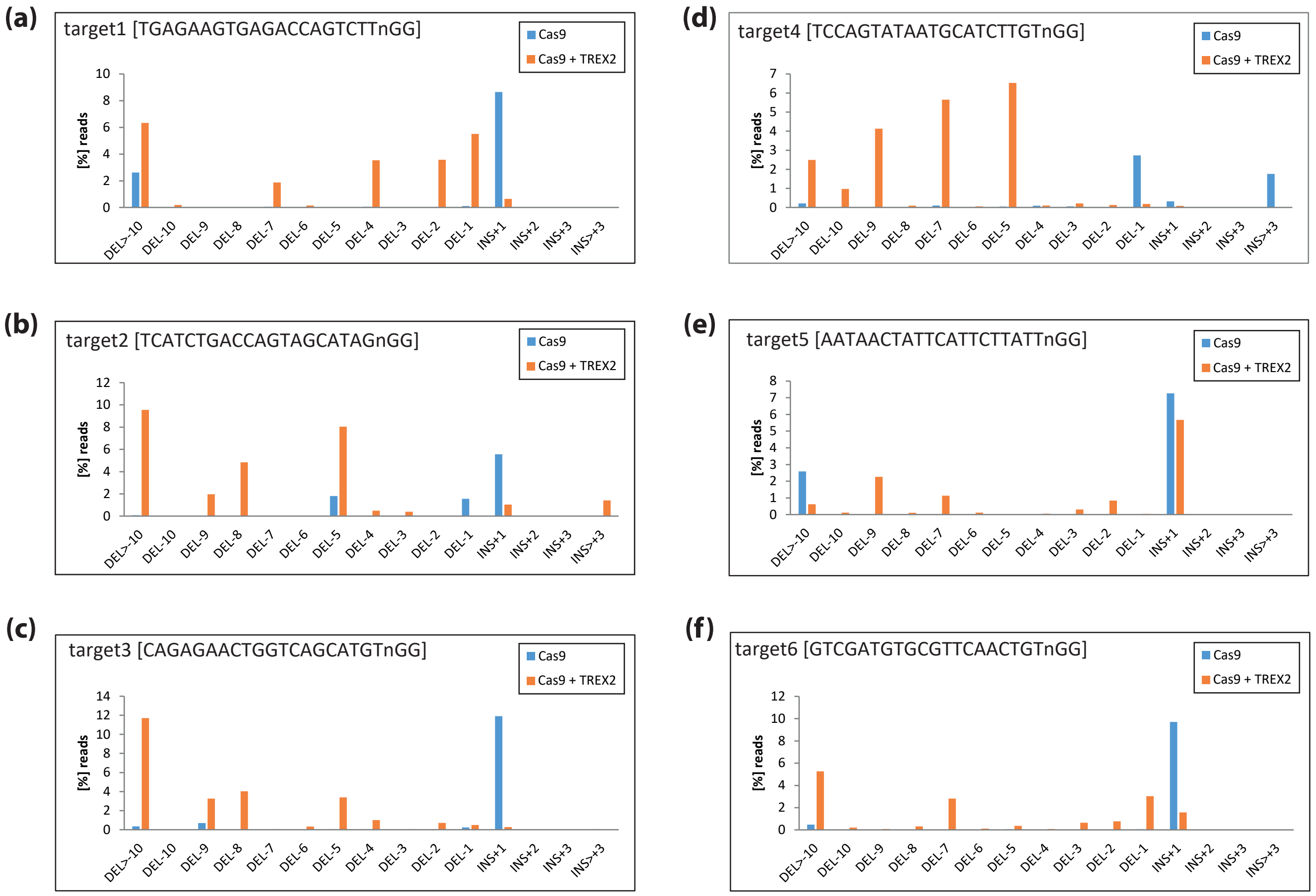


**Figure S5:** Mutation (InDel) profiles in absence/presence of TREX2 at single target sites.

Panels a)-f) show InDel profiles as determined by amplicon sequencing and CRISPresso analysis for individual target sites. For example, graphs a) and b) represent data obtained using DNA from $T_1$ transformants from pDGE1081 and pDGE1082.
