## Supplementary material for "Targeted gene deletion with *Sp*Cas9 and multiple guide RNAs in *Arabidopsis thaliana*: four are better than two": Figure S6

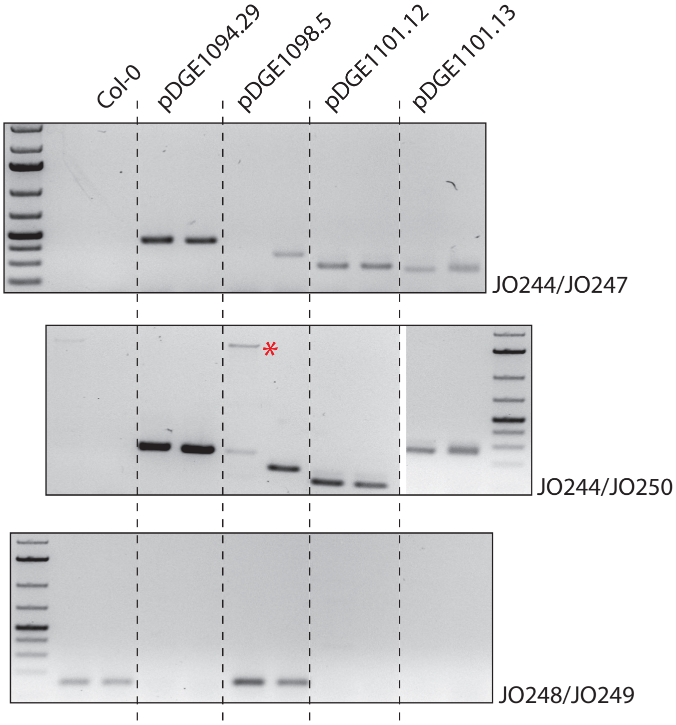


**Figure S6:** PCR-genotyping of putative *wrky30* deletion lines in the $T_2$ generation.

Non-transgenic seeds were selected from indicated $T_2$ populations (see Figure S4 for genotyping in the $T_1$ generation) using the FAST marker. Four plants were pooled for DNA extraction; two pools per population were genotyped by PCR for presence/absence (deletion) of the *WRKY30* locus. Amplicons JO244/247 and JO244/250 both query deletion of *WRKY30* (see Figure 1 for primer binding sites). A product corresponding to the length of the wild-type *WRKY30* locus was amplified from one pool DNA derived from 1098.5 (marked with an asterisk). Amplicon JO248/249 queries presence of the *WRKY30* locus. PCR results confirm bi-allic deletions at the *WRKY30* locus (Figure S4) for populations 1094.29, 1101.12 and 1101.13.
