## Supplemental File S2 for "Targeted gene deletion with *Sp*Cas9 and multiple guide RNAs in *Arabidopsis thaliana*: four are better than two"

**Supplemental File S2: Plasmids, oligonucleotides, cloning details**

**Previously published plasmids:**

| name | Description | Reference |
|---|---|---|
| pDGE331 | M1E sgRNA shuttle | (Stuttmann *et al.*, 2021) |
| pDGE332 | M1 sgRNA shuttle | (Stuttmann *et al.*, 2021) |
| pDGE333 | M2 sgRNA shuttle | (Stuttmann *et al.*, 2021) |
| pDGE334 | M2E sgRNA shuttle | (Stuttmann *et al.*, 2021) |
| pDGE335 | M3 sgRNA shuttle | (Stuttmann *et al.*, 2021) |
| pDGE336 | M4 sgRNA shuttle | (Stuttmann *et al.*, 2021) |
| pDGE337 | M4E sgRNA shuttle | (Stuttmann *et al.*, 2021) |
| pDGE495 | M5 sgRNA shuttle | (Stuttmann *et al.*, 2021) |
| pDGE497 | M6E sgRNA shuttle | (Stuttmann *et al.*, 2021) |
| pCK70 | zCas9i(2xNLS) (pAGM47523) [CDS1] | (Stuttmann *et al.*, 2021)<br>(Grützner *et al.*, 2021) |
| pCK73 | p2x35S:zCas9i-tRbcSE9 [1-2f] | (Stuttmann *et al.*, 2021) |
| pCK75 | pBs3:Bs3-tocs [1-3f] | (Stuttmann *et al.*, 2021) |
| pCK76 | pnos:nptII-tnos [1-5f] | (Stuttmann *et al.*, 2021) |
| pTEI44 | pRPS5a [Pro+5U(f)] | (Gantner *et al.*, 2018) |
| pJOG292 | Acceptor for pDGE recipient assembly | (Ordon *et al.*, 2017) |
| pJOG294 | ccdB cassette for pDGE recipient assembly | (Ordon *et al.*, 2017) |
| pJOG416 | trbcS-E9 [3U+ter] | (Gantner *et al.*, 2018) |
| pJOG603 | pRPS5a [Pro+5U] | (Gantner *et al.*, 2018) |
| pJOG640 | p35S [Pro+5U(f)] | (Gantner *et al.*, 2018) |
| pJOG648 | pAt2S3 [Pro+5U] | (Gantner *et al.*, 2018) |
| pJOG990 | NbEU terminator [3U+Ter] | (Stuttmann *et al.*, 2021)<br>(Diamos and Mason, 2018) |
| pJOG1008 | t35S::tNbAct::Rb7MAR terminator (ttriple) [3U+Ter] | (Diamos and Mason, 2018,<br>Stuttmann *et al.*, 2021) |
| pICH47761 | Level 1 acceptor | (Engler *et al.*, 2014) |
| pICH47742 | Level 1 acceptor | (Engler *et al.*, 2014) |
| pICH47751 | Level 1 acceptor | (Engler *et al.*, 2014) |
| pICH41800 | End-linker | (Engler *et al.*, 2014) |
| pICH41766 | End-linker | (Engler *et al.*, 2014) |
| pICHSL80007 | mCherry [CDS1] | (Engler *et al.*, 2014) |
| pICH72400 | tug7 [3U+ter] | (Engler *et al.*, 2014) |
| pICH51288 | 2x35S [Pro+5U] | (Engler *et al.*, 2014) |
| pICSL70008 | FAST [gene] | (Engler *et al.*, 2014) |

**Level 0 modules (cloned as described in Engler at al., 2014):**

| name | recipient | Description | oligonucleotides |
|---|---|---|---|
| pJOG983 | pGAM1276 | TREX2-P2A as NT1 | JS1684/1685 on plasmid Addgene #91026 (Cermak et al., 2017) |

**Level 1 modules (assembled as described in Engler et al., 2014):**

| name | recipient | Inserts | Description |
|---|---|---|---|
| pJOG685 | pICH47761 | pJOG648, pICHSL80007, pICH72400 | pAtS2S3:mCherry-tug7 [1-4f] |
| pJOG991 | pICH47742 | pJOG640, pJOG983, pJOG416, pCK70 | p35S:TREX-zCas9i-trbcS [1-2f] |
| pJOG1030 | pICH47742 | pICH51288, pCK70, pJOG990 | p35S:zCas9i-tNbEU [1-2f] |
| pJOG1031 | pICH47742 | pICH51288, pCK70, pJOG1008 | p35S:zCas9i-ttriple [1-2f] |
| pJOG304 | pICH47751 | pICSL70008 | FAST [1-3f] |
| pCK226 | pICH47742 | pJOG603, pCK70, pJOG1008 | pRPS5a:zCas9i_ttriple [1-2f] |
| pCK227 | pICH47742 | pTEI44, pJOG983, pCK70, pJOG1008 | pRPS5a:TREX2-zCas9i_ttriple [1-2f] |

**pDGE – recipients (assembled as described for Level 2 constructs in Engler et al., 2014):**

| name | recipient | inserts | description |
|---|---|---|---|
| pDGE1108 | pJOG292 | pJOG304, pCK226, pJOG294, pICH41766 | FAST_ pRPS5a:Cas9(ttriple)_ccdB |
| pDGE1109 | pJOG292 | pJOG304, pCK227, pJOG294, pICH41766 | FAST_ pRPS5a:TREX-Cas9(ttriple)_ccdB |
| pDGE311 | pJOG292 | pCK73, pCK75, pCK76, pJOG685, pJOG294, pICH41800 | nptII-Bs3-Cherry-35S:Cas9_rbcS-ccdB |
| pDGE345 | pDGE311 | pDGE331 (empty M1E module) | nptII-Bs3-Cherry-35S:Cas9_rbcS-M1E |
| pDGE355 | pJOG292 | pJOG294, pJOG685, pJOG991, pCK75, pCK76, pICH41800 | nptII-Bs3-Cherry-35S:TREX2-Cas9_rbcS-ccdB |
| pDGE399 | pJOG292 | pJOG1030, pCK75, pCK76, pJOG685, pJOG294, pICH41800 | nptII-Bs3-Cherry-35S:Cas9_NbEU-ccdB |
| pDGE400 | pJOG292 | pJOG1031, pCK75, pCK76, pJOG685, pJOG294, pICH41800 | nptII-Bs3-Cherry-35S:Cas9_ttriple-ccdB |

**pDGE - sgRNA shuttle vectors (constructed as described in Stuttmann et al., 2021):**

| name | recipient | Oligonucleotides | Module type |
|---|---|---|---|
| pDGE390 | pDGE331 | JS809/810 | M1E |
| pDGE1079 | pDGE332 | JS2642/2643 | M1 |
| pDGE1080 | pDGE334 | JS2648/2649 | M2E |
| pDGE1083 | pDGE332 | JS2644/2645 | M1 |
| pDGE1084 | pDGE334 | JS2646/2647 | M2E |
| pDGE1087 | pDGE332 | JS2640/2641 | M1 |
| pDGE1088 | pDGE334 | JS2650/2651 | M2E |
| pDGE1091 | pDGE333 | JS2644/2645 | M2 |
| pDGE1092 | pDGE335 | JS2648/2649 | M3 |
| pDGE1093 | pDGE337 | JS2646/2647 | M4E |
| pDGE1096 | pDGE333 | JS2640/2641 | M2 |
| pDGE1097 | pDGE337 | JS2650/2651 | M4E |
| pDGE1100 | pDGE335 | JS2650/2651 | M3 |
| pDGE1103 | pDGE336 | JS2640/2641 | M4 |
| pDGE1104 | pDGE495 | JS2646/2647 | M5 |
| pDGE1105 | pDGE497 | JS2650/2651 | M6E |

**pDGE – editing constructs (assembled as described in Stuttmann et al., 2021):**

| name | recipient | inserts | description |
|---|---|---|---|
| pDGE375 | pDGE345 | Oligonucleotides JS809/810 | nptII-Bs3-Cherry-35S:Cas9_rbcS-sgRNA |
| pDGE404 | pDGE355 | pDGE390 (sgRNA vs. NbEDS1) | nptII-Bs3-Cherry-35S:TREX2-Cas9_rbcS-sgRNA |
| pDGE405 | pDGE399 | pDGE390 (sgRNA vs. NbEDS1) | nptII-Bs3-Cherry-35S:Cas9_NbEU-sgRNA |
| pDGE406 | pDGE400 | pDGE390 (sgRNA vs. NbEDS1) | nptII-Bs3-Cherry-35S:Cas9_ttriple-sgRNA |
| pDGE1081 | pDGE1108 | pDGE1079, 1080 | WRKY editing 2 sgRNAs |
| pDGE1082 | pDGE1109 | pDGE1079, 1080 | WRKY editing 2 sgRNAs TREX |
| pDGE1085 | pDGE1108 | pDGE1083, 1084 | WRKY editing 2 sgRNAs |
| pDGE1086 | pDGE1109 | pDGE1083, 1084 | WRKY editing 2 sgRNAs TREX |
| pDGE1089 | pDGE1108 | pDGE1087, 1088 | WRKY editing 2 sgRNAs |
| pDGE1090 | pDGE1109 | pDGE1087, 1088 | WRKY editing 2 sgRNAs TREX |
| pDGE1094 | pDGE1108 | pDGE1079, 1091,1092,1093 | WRKY editing 4 sgRNAs |
| pDGE1095 | pDGE1109 | pDGE1079, 1091,1092,1093 | WRKY editing 4 sgRNAs TREX |
| pDGE1098 | pDGE1108 | pDGE1079,1096,1092, 1097 | WRKY editing 4 sgRNAs |
| pDGE1099 | pDGE1109 | pDGE1079,1096,1092, 1097 | WRKY editing 4 sgRNAs TREX |
| pDGE1101 | pDGE1108 | pDGE1083, 1096, 1100,1093 | WRKY editing 4 sgRNAs |
| pDGE1102 | pDGE1109 | pDGE1083, 1096, 1100,1093 | WRKY editing 4 sgRNAs TREX |
| pDGE1106 | pDGE1108 | pDGE1079,1091,1092, 1103,1104,1105 | WRKY editing 6 sgRNAs |
| pDGE1107 | pDGE1109 | pDGE1079,1091,1092, 1103,1104,1105 | WRKY editing 6 sgRNAs TREX |

**Oligonucleotides used for cloning of Level 0 modules:**

| name | sequence |
|---|---|
| JS1684 | ttgaagacatCCATGTCTGAGCCACCTCGGGCTGAaACCTTTG |
| JS1685 | ttgaagacatCATtGGTCCAGGATTTTCTTCAAC |

**Oligonucleotides for sgRNA construction:**

| name | sequence |
|---|---|
| JS809 | ATTGTATGCTGCATGTAATCTGAA |
| JS810 | AAACTTCAGATTACATGCAGCATA |
| JS2640 | attgAATAACTATTCATTCTTATT |
| JS2641 | aaacAATAAGAATGAATAGTTATT |
| JS2642 | attgTGAGAAGTGAGACCAGTCTT |
| JS2643 | aaacAAGACTGGTCTCACTTCTCA |
| JS2644 | attgCAGAGAACTGGTCAGCATGT |
| JS2645 | aaacACATGCTGACCAGTTCTCTG |
| JS2646 | attgTCCAGTATAATGCATCTTGT |
| JS2647 | aaacACAAGATGCATTATACTGGA |
| JS2648 | attgTCATCTGACCAGTAGCATAG |
| JS2649 | aaacCTATGCTACTGGTCAGATGA |
| JS2650 | attGTCGATGTGCGTTCAACTGT |
| JS2651 | aaacACAGTTGAACGCACATCGA |

**Oligonucleotides used for genotyping, sequencing:**

| name | Sequence [purpose] | |
|---|---|---|
| JS1753 | GCGATCAGATTCTCAAGCCG | [zCas9i presence/absence] |
| JS1754 | TTTTGCAGGTTGACGACTCG | [zCas9i presence/absence] |
| JO244 | TTGTCTCTTGGAATTTCTAAACTCAA | [WRKY30 genotyping] |
| JO245 | TTTGACTGAAGAACGAAGAAAGCT | [WRKY30 genotyping] |
| JO246 | TGCAAATTTGAGTCTTCTTTTAGCT | [WRKY30 genotyping] |
| JO247 | TCTGTGGTAGAGAAATTAAAGAGGT | [WRKY30 genotyping] |
| JO248 | CCACTCTTTGAACGTAATGGAGAA | [WRKY30 genotyping] |
| JO249 | TTTGGCTAAATGTTCACGTGTTTC | [WRKY30 genotyping] |
| JO250 | AGAAAAGTTTATCTGTCTGTGGTAGA | [WRKY30 genotyping] |
| JS1132 | AACGCTCTTTTCTCTTAGGT | [sgRNA array sequencing] |
| JS2302 | GTAATAGCAATGACCAGTGC | [sgRNA array sequencing] |

### References


**Diamos, A.G. and Mason, H.S.** (2018) Chimeric 3' flanking regions strongly enhance gene expression in plants. *Plant Biotechnol J*.

**Engler, C., Youles, M., Gruetzner, R., Ehnert, T.M., Werner, S., Jones, J.D., Patron, N.J. and Marillonnet, S.** (2014) A Golden Gate Modular Cloning Toolbox for Plants. *ACS synthetic biology*.

**Gantner, J., Ordon, J., Ilse, T., Kretschmer, C., Gruetzner, R., Lofke, C., Dagdas, Y., Burstenbinder, K., Marillonnet, S. and Stuttmann, J.** (2018) Peripheral infrastructure vectors and an extended set of plant parts for the Modular Cloning system. *PLoS ONE*, **13**, e0197185.

**Grützner, R., Martin, P., Horn, C., Mortensen, S., Cram, E.J., Lee-Parsons, C.W.T., Stuttmann, J. and Marillonnet, S.** (2021) High-efficiency genome editing in plants mediated by a Cas9 gene containing multiple introns. *Plant Communications*, **2**, 100135.

**Ordon, J., Gantner, J., Kemna, J., Schwalgun, L., Reschke, M., Streubel, J., Boch, J. and Stuttmann, J.** (2017) Generation of chromosomal deletions in dicotyledonous plants employing a user-friendly genome editing toolkit. *Plant J*, **89**, 155-168.

**Stuttmann, J., Barthel, K., Martin, P., Ordon, J., Erickson, J.L., Herr, R., Ferik, F., Kretschmer, C., Berner, T., Keilwagen, J., Marillonnet, S. and Bonas, U.** (2021) Highly efficient multiplex editing: one-shot generation of 8x Nicotiana benthamiana and 12x Arabidopsis mutants. *Plant J*, **106**, 8-22.
